# Airway smooth muscle--on-a-chip: a microfluidic approach to study alveolar smooth muscle remodelling

**DOI:** 10.1101/2025.03.13.643111

**Authors:** Amir Suhail, Joseph Xavier, PK Hiba, Ajai Krishnan MJ, Amritha Pradeep, KB Megha, S Reshma, NS Remya, Jorge Bernardino de la Serna

## Abstract

Respiratory illnesses, like chronic obstructive pulmonary disease (COPD) and asthma, pose significant global health challenges due to their chronic nature and limited treatment options. Airway smooth muscle (ASM) plays a vital role in respiratory diseases, particularly in airway remodelling and obstruction. ASM, which encircles the bronchial tree and extends to the trachea, plays a vital yet not fully understood role in lung physiology. However, its dysfunction is strongly associated with asthma and COPD progression, leading to excessive contraction, increased inflammatory mediator release, and ASM hypertrophy. However, identifying its precise function is challenging due to limitations in existing research models for assessing ASM contraction. In vivo models offer a comprehensive physiological perspective but possess ethical concerns and they do not allow for the direct measurement of ASM contraction. Meanwhile, ex vivo and in vitro models provide a more direct assessment; however, they lack crucial physiological factors. Understanding how ASM cells interact with their surroundings is essential for gaining deeper insights into respiratory disorders. To address this gap, we aimed to mimic the human airway smooth muscle-on-a-chip model, incorporating ASM cells in a 3D microenvironment. This microfluidic platform provides a physiologically relevant environment, allowing for studying complex mechanisms that drive airway remodelling and dysfunction in respiratory diseases. The ASM-on-a-chip is designed for long-term 3D cell culture of ASM cells that reorient itself to form a smooth muscle fibre. The design provides side channels for manipulating the constituent of the hydrogel to study the effect of compounds on AMS remodelling.

## Introduction

According to the World Health Organisation (WHO) 99% of the world’s population breath polluted air (above the WHO guideline limit). ^1^ Global climate change has taken the release of air pollutants on an exponential curve. Alarming increases in extreme weather events, intense wildfires, floods, heat waves and hurricanes directly increase the risk of exposure to hazardous particles which has aggravated respiratory disorders in the world population. ^2,3^ Inhaling hazardous materials and polluted air triggers chronic inflammation, hypoxia, and the generation of Reactive Oxygen Species (ROS), activating the immune response. ^4^ This, in turn, increases the local production of inflammatory markers and growth factors. ^5^ The continuous release of these substances disrupts tissue repair mechanisms and alters airway structure, ultimately leading to pathological conditions. ^6,7^

Respiratory disorders like asthma and chronic obstructive pulmonary disease (COPD) are characterised by structural changes in the airways. Airway wall thickening in such conditions significantly impacts lung function. Airway smooth muscle (ASM) is a crucial tissue responsible for controlling broncho-motor tone and is found in the trachea as well as throughout the bronchial tree, extending to the terminal bronchioles. ^8^ ASM cells play a key role in airway function by facilitating contraction. Changes in ASM cell function, including altered contractility, are closely associated with airway inflammation, hyperresponsiveness, and structural remodelling which drastically hinders respiration mechanics reducing the airflow, eventually exacerbating disease severity. ^9^

Moreover, ASM remodelling plays a significant role in the progression of asthma and COPD. In asthma, increased ASM mass contributes to airway narrowing and hyperresponsiveness. This process involves ASM hypertrophy and hyperplasia, leading to thickening of the airway walls. Additionally, ASM cells produce pro-inflammatory cytokines and extracellular matrix proteins, promoting airway inflammation and fibrosis. These structural changes result in airflow obstruction and respiratory dysfunction. ^10^ In COPD, airway smooth muscle (ASM) dysfunction affects the contractility, immune responses, and airway structure, leading to remodelling and fibrosis. These alterations impact lung function and elasticity. ^11^ Cytokines like interleukin-5 (IL-5), transforming growth factor-beta (TGF-β), and tumour necrosis factor-alpha (TNF-α) act as key regulators of ASM remodelling. IL-5 plays a role in activating and maintaining eosinophils and mast cells, which are crucial in allergic asthma. TGF-β and TNF-α promote ASM cell growth, thickening, and the production of extracellular matrix (ECM) proteins, leading to airway structural changes. ^10^ ASM cells are a source of ECM proteins that drive structural changes in the airway. The interaction between ASM cells and the ECM can influence ASM function and contribute to airway stiffening. ^12^ Understanding the mechanisms underlying ASM remodelling in these diseases is crucial for developing targeted therapies aimed at mitigating airway remodelling and improving patient outcomes.

Organs-on-chips are microfluidic platforms designed to grow living cells in small, perfused chambers, mimicking the physiological functions of tissues and organs. ^13^ To understand human health and diseases, we need to learn how cells work in their natural tissue and organ microenvironments. ^14^ While research has identified key factors such as extracellular matrix modifications, growth factors, and mechanical forces, but how these elements interact dynamically within the lung microenvironment is yet to be understood. ^15^ An important study by Humayun et al 2018, created a lung-on-a-chip device to mimic the interface between primary airway epithelial cells (ECs) and smooth muscle cells (SMCs), in an ECM. ^16^ However, the gap in airway smooth muscle (ASM) remodelling studies lies in the limited understanding of the molecular mechanisms driving structural changes in pathological conditions like asthma and COPD. Additionally, most studies still rely on in vitro or animal models, which may not fully capture human disease complexity. ^17^ To fullfill this gap and enable dynamic ASM studies in detail we developed a microfluidic ASM-on-chip that can mimic three-dimensional ASM microenvironments and recapitulate ASM remodelling. This would allow us to understand the triggering factors of ASM remodelling and aid us in investigating treatment possibilities. The ASM-on-chip stands out for its ability to support the three-dimensional culture of Primary Lung Smooth Muscle (PLSM) cells, allowing them to structurally reorient into a single muscle fibre within the chip. The design of our ASM-on-chip enables the exposure of potentially hazardous molecules and inflammatory signals for toxicity and functional analysis, as well as testing potential therapeutics to reverse ASM remodelling.

## Experimental

### Cultivation of cells and hydrogel optimisation

Primary Lung Smooth Muscle (PLSM) cells were purchased from the American Type Culture Collection (ATCC, PCS-130-010) and cultured on T25 culture flasks in Nutrient mixture F-12 Ham, with Kaighn’s modification (F12 Ham’s media) (Himedia, AT106-10X) supplemented with 10% FBS (Gibco, 2585261) and 1% Antibiotic antimycotic solution (A002, Himedia). Similarly, Adenocarcinomic human alveolar basal epithelial cells (A549) were purchased from NCCS (Pune, India) and maintained in MEM nutrient medium (ALAT154, Himedia) supplemented with 10% FBS and 1% Antibiotic antimycotic solution (A002, Himedia). Cells were maintained in a humidified 37C incubator and media were replaced every two days. Cells at passages 5 to 8, after attaining 80% confluency cells were detached from the flask using Trypsin-EDTA solution (Himedia, TCL070) and used for the experiments. To mimic the three-dimensional environment of PLSM cells inside the chip collagen 1, rat tail (Gibco, A10483-01) was used. Prior to microfluidic cell culture, the optimal collagen concentration was determined by seeding PLSM cells in different concentrations of collagen 1. Briefly, three different concentrations were used: 1 mg/mL, 1.5 mg/mL, and 2 mg/mL. The hydrogel was prepared by mixing Collagen I, 10X F12 Ham’s media,1 Normal NaOH, Sterilized water and cell suspension as per the manufacturer’s protocol. The pH was adjusted between 6.5 to 7.5, and finally, the cell suspension was added to adjust the final concentration to 105 cells per mL. The mixture was seeded into a 96-well plate and placed in a humidified incubator at 37°C for 30 to 40 minutes to set the gel following the addition of 50 µL F12 Ham’s media. After 24 hours of incubation, the cell was observed under phase contrast microscopy (Olympus CKX53).

### Live/dead assay

A live/dead assay for gel optimization using calcein-AM/propidium iodide was performed to evaluate the cell proliferation and visualise the cell within the collagen hydrogel. This will help to determine the hydrogel concentration best suited for the 3D culture of PLSM that suspends the cells evenly throughout the hydrogel. Membrane permeable calcein-AM is cleaved by intracellular esterase in the live cells to form green, fluorescent calcein molecule in the cytoplasm.. The calcein-AM stain (2µM) was added into the wells and incubated at 37°C for 15 min, after removing the stain from the wells, it was washed with 1X PBS, 2 times. PI (4 µM) was added and incubated at 37°C for 10 min. PI is an impermeable red, fluorescent dye penetrating the cell when it loses its plasma membrane integrity. The wells were then washed using 1X PBS and the cells were observed under confocal laser scanning microscopy (CLSM) (Olympus IX83, Japan) using 10x objective (NA 0.3). 3D stacked images of each well (step size 5µm) were taken by selecting FITC and PI dyes; excited with 488nm and 561nm solid state lasers and the emission was collected through 500-540nm and 570-670nm bandpass filters for the respective dyes. Later the images were analysed for uniformity of cell distribution to fix the collagen 1 concentration.

### Design and fabrication of the chip

The microfluidic device consists of three parallel channels, each with a width of 1000 μm and a height of 200 μm, separated by 100 μm apart micropillars with a diameter of 250 μm (Fig S1). This design was based on previous literature ^18^ and modified to optimise for this study. The central channel is designated for hydrogel seeding, and the side channels are used for media filling. The micropillar design enables the diffusion of chemicals and nutrients across the gel channel from the media channel.

Once the design of the device was finalised, the positive mould for the ASM-on-a-chip was drawn using AutoCAD software. G-code was developed for each toolpaths and sent to the Wegstr CNC controlling software (Wegstr, Czech Republic). A polymethylmethacrylate (PMMA) sheet, 6 mm in depth (50 × 75 mm width x height), was secured on the Wegstr stage using double-sided tape. For milling the micropillars, a 200 μm drill bit (Wegstr, Czech Republic) was used, while the channel borders were milled using a 300 μm end mill (Wegstr, Czech Republic). The PMMA mould was released from the CNC stage, cleaned using isopropanol in an ultrasonic bath (2 minutes), and dried using nitrogen gas. The cleaned mould was exposed to chloroform vapour for five minutes and kept in a hot-air oven at 60°C for 2 hours to improve its surface smoothness. Furthermore, the chip mould was exposed to 1H,1H,2H,2H-Perfluoro-octyl-triethoxysilane (667420, Sigma Aldrich) to render the mould surface more hydrophobic and to improve the peeling of cured PDMS from the mould.

Sylgard 184 PDMS (Dow Chemicals) oligomer is mixed with its curing agent in a 10:1 weight ratio and degassed using a vacuum desiccator to remove any air bubbles trapped during mixing. The degassed PDMS mixture is poured into the PMMA mould and degassed again to remove any bubbles trapped in the micropillars. Subsequently, the mould is placed in a hot-air oven at 60°C for 2 hours to cure the PDMS. The cured PDMS is peeled off from the PMMA mould, and the edges are trimmed. The inlets and outlets are punched using a biopsy puncture (2mm and 5mm for central and side channels; Netcare instruments, India). The PDMS chips and glass coverslips (25 × 60 mm) are cleaned in a bath sonicator and dried in a hot-air oven. The cleaned PDMS chips and glass coverslips are exposed to air plasma (Harrick Plasma, USA) for two minutes (high RF power). The plasma-exposed PDMS chips and glass slides are bonded together in conformal contact and kept in a hot-air oven at 60°C for 2-3 hours. The chips are then sterilized by autoclave and dried prior to use.

### Numerical simulation

In this study, a numerical model was developed using COMSOL Multiphysics (COMSOL, Inc.) to simulate the gel-filling process of the collagen (2mg/mL) through the central channel of the chip. Then, it was validated by passing blue-coloured water through the central channel and by injecting the collagen hydrogel. The present microfluidic chip was designed based on the capillary burst valve model ^19^. The uniformly spaced circular micropillars act as geometric capillary burst valves. The interface of the fluids gets pinned in between the gaps of the pillars. A partial 2-D model of the current design with two space dimensions (collagen and air), 23 domains and 90 boundaries were uploaded into the computational fluid dynamics (CFD) software to simulate and understand the gel filling inside the central channel. The CFD simulation integrated the incompressible Navier-Stokes equation, the continuity equation, and the two-phase level set method ^20^.

The primary parameters influencing the gel-filling process encompass the viscosity and surface tension coefficient of the mixture, the velocity at which the gel is filled, the contact angle of the microchannel surfaces, and the spacing between the micropillars ^19^. The concentration of the collagen was 2 mg/mL, which was set earlier as per the cell distribution; the density of collagen solution (2 mg/ml) was set at 1000 kg/m^3^, the dynamic viscosity was set at 6mPa.s, and the surface tension of the gel channel was 0.07 N/m ^20^. A wetted wall boundary condition was implemented on the inner walls of the central channel. Thus, the contact angle between the liquid interface and the polydimethylsiloxane (PDMS) side wall was set to 140*π/180, which plays a pivotal role in the gel filling ^20^. The collagen filling flow rate was set to 30 µL/min. A time-dependent simulation of 0 to 1 sec was conducted with output intervals of 0.1 sec. The numerical simulation underwent meshing with various sizes of free triangular meshes. The accuracy of the collagen solution filling was assessed by employing a grid convergence study. A physics-controlled meshing method was used to mesh the geometry to increase the mesh resolution near the boundaries and corners of the fluidic domain, thus reducing the errors in the simulation.

### ASM-on-a-chip cell culture and characterization

The fabricated chips were autoclaved before starting cell culture to ensure the sterility of the chips. For hydrogel preparation, the optimal concentration of collagen 1 fixed from the gel optimisation study was used. PLSM cells after attaining 80% confluency were trypsinized from a T25 flask. 2 mg/ml Collagen 1 hydrogel was prepared by mixing Collagen 1, 10X F-12 ham’s K media, 1N NaOH, cell suspension (adjusted to a final concentration of 5.1×10^5^ cell/mL, and sterile water to make up the solution into 400 µL (the total volume can be changed based on number chips). The hydrogel was mixed thoroughly to ensure an even distribution of cells in the mixture. 10 µL of hydrogel was seeded carefully into the sterile chip via the central inlet, till the central channel was filled. The chip was then incubated at 37°C for 30 min for gelation. Following gelation, culture media was added through both side channels to supplement the cells present in the hydrogel with nutrients. Media in the side channels were changed every 24 hours to ensure viability for 7 days. Cells were observed daily and imaged under a phase contrast microscope. For characterising the long-term culture of PLSM cells using the chip, live/dead assay using calcein-AM and PI staining and F-actin staining using rhodamine-phalloidin was done (2 days, 7 days after cell seeding). Similarly, A549 cells in collagen hydrogel were also loaded in the chip to investigate cell-dependent reorganization and was maintained for 2 days. Live/dead assay is performed as briefed above, except that the stains and PBS for washing steps were added via the side channels.

We image the cells by immune staining using protocols described before. ^21^ Similarly, F-actin was stained using Rhodamine phalloidin (Ab176756) and counter-stained with 4,6 Diamidino-2-phenylindole (DAPI). It helps to evaluate the shape, orientation, and organization of the cells. The chips from days 2 and 7 were fixed using 4% paraformaldehyde for 10 minutes in the dark. Following fixation, the cells were washed with 1X PBS, 3 times. For permeabilization, 0.2% Triton X-100 was added for 7 minutes, followed by 1X PBS wash 3 times. DAPI (1 µg/mL) was added and incubated at room temperature for 5 min, followed by 1X PBS wash 3 times. Rhodamine phalloidin (1x) was then added and incubated at room temperature for 10 min. Chips were washed again with 1X PBS, and the cells were observed under a confocal laser scanning microscope. 3D stacked images were taken using 10x objective lens (NA 0.3), selecting rhodamine phalloidin and DAPI dyes; excited with 561nm and 405nm solid state lasers and the emission was collected through 570-670nm and 430-470nm bandpass filters for the respective dyes.

Fluorescein Isothiocyanate (FITC) conjugated Dextran-70 kDa (Sigma Aldrich, USA) was used to study the diffusion of molecules across the collagen hydrogel. The ASM chip was loaded with collagen 1 hydrogel without the cells and allowed gelation for 30 min at 37°C. 100 µL 1mM FITC-Dextran was added into the side channels under the confocal microscope and images were taken every 5 mins for 30 mins.

### TGF-β treatment in ASM-on-a-chip

After seeding the cells in the chip (day -2) the cells were supplemented with ham’s F-12 K media with 10% FBS for 24 hours. Then cells were cultured under serum-deprived condition (ham’s F-12 K media with 0.5% FBS) for the next 24 hours before starting the treatment. At day 0 (treatment), chips were divided into two groups, TGF-β – (media without TGF-β) and TGF-β + (media with 10 ng/mL TGF-β (CF078, Himedia, India)). Media was refreshed every day for 7 days and images were taken daily to observe the changes in gel compaction and fibre formation. On days 2 and 7 chips were fixed using 4% paraformaldehyde for staining and further analysis.

### Immunofluorescence staining and confocal imaging

On the 2nd and 7th days of TGF-β treatment, immunocytochemistry was done to analyse the expression of calponin-1, a contractile marker expressed by PLSM cells. The culture medium was removed from the chips and the cells were fixed by adding 4% paraformaldehyde (PFA) (Sigma Aldrich, USA) to the side channels, followed by a 15-minute incubation at room temperature. After removing the PFA, the chips were washed with 1X PBS 3 times. The cells were then permeabilized with 0.25% Triton X-100, incubated for 10 minutes at room temperature and washed with 1X PBS. 200 µL of blocking buffer (1% bovine serum albumin (BSA) (Sigma Aldrich, USA) and 22.52 mg/mL glycine (SRL, India) in PBST) were added to the side channels followed by a 30 min incubation at room temperature. Then, 200 µL of Anti-calponin 1 antibody [EP798Y] (Cat. No: ab46794) was diluted to 1:100 in 1% BSA in PBST and added to incubate overnight at 4°C in a humidified chamber, the primary antibody was removed and washed with 1X PBS. 100 µL of secondary antibody Goat Anti-Rabbit IgG H&L (Alexa Fluor-488, (Cat. No: ab150077) was diluted 1:400 in 1% BSA in PBST was added to the chip and incubated for 2 hours in the dark at room temperature. The secondary antibody was washed with 1X PBS and counterstained with DAPI (1 µg/mL) for 5min. The chips were washed again with 1X PBS before acquiring images under a Confocal laser scanning microscope. 3D stacked images were taken using 10x (NA 0.3) and 20x (NA 0.8) objective lens, selecting Alexa Fluor-488 and DAPI dyes; excited with 488nm and 405nm solid state lasers and the emission was collected through 500-600nm and 430-470nm bandpass filters for the respective dyes.

### Image analysis

#### Hydrogel contraction

The phase contrast images taken over 7 days were analysed using Olympus cellSens standard software. Briefly, triplicate images of TGF-β – and TGF-β + groups were taken from day 0 to day 7. Using an arbitrary line measurement tool, widths at multiple sections were measured from each image. The data was then processed, and percentage contraction was calculated using the following equation:

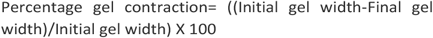

### Cell counts and nuclear orientation

To quantify and compare the number of cells in TGF-β – and TGF-β + groups at days 2 and 7, the number of nuclei was quantified from Z-stacked images of Rhodamine-DAPI staining. Image analysis software (Fiji ImageJ 1.54g) was used to perform image cytometry. Briefly, the red (F-actin) and blue channels (Nuclei) of the images were split, and the blue channel was stacked for maximum intensity projection over the z-axis. A Gaussian blur was applied to smoothen the nuclei, and the threshold was adjusted to separate nuclei from each other. An ImageJ plug-in called MorphoLibJ was used for segmentation and automatic counting of the nuclei.^22^ Values acquired from the table were then exported to Microsoft Excel for analysis.

The orientation of the cell in 3D hydrogel can be determined by analysing the orientation of the nuclei in DAPI-stained images. The alignment of cells between day 2 and day 7 of TGF-β – chips were compared to evaluate the time-dependent effect of PLSM orientation inside the chip. Similar to the protocol of cell counting, the ‘Equivalent Ellipse’ analysis algorithm in the MorphoLibJ plug-in of ImageJ software is used to obtain nucleus orientation. Origin Pro 8.5 software was used to plot the frequencies of bin orientation angles as a rose diagram.

### Fluorescent intensity measurements

To quantify the expression level of F-actin and Calponin from Z-stacked images of Rhodamine-DAPI staining and anti-calponin staining, Fiji ImageJ software was used. Measurements of mean grey value and integrated density were set in the “Set measurement” menu and data were gathered from the “Measure stack” to acquire intensity values from each stack. For the analysis of calponin expression, equal number of stacks were taken from all groups, and the sum of integrated density was normalised by dividing the number of cells in both groups.

### Statistical analysis

Data plotted as graphical representations are interpreted as mean value ± SD. A minimum of 3 replicates were taken for analysis. Unpaired t-test was used to compare gel contraction between the groups. Two-way ANOVA and Sidak’s multiple comparison test were used to compare the fluorescence intensity and cell number between the groups. Statistical significance was defined as p < 0.05 (*), p < 0.01 (**), p < 0.0001 (****) and p > 0.05 as non-significant (ns).

## Results and discussion

### PLSM culture and hydrogel optimisation

Primary Lung Smooth Muscle cells attaining confluency in culture flask exhibit typical hill and valley patterns. Cell exhibits behavioural and expression changes when cultured in 3D environments compared to 2D cultures owing to the differences in the cell-extracellular matrix (ECM) and cell interactions. Furthermore, complex ECM constituents and biophysical stimuli imparted via ECM activate and control cell behaviours in vivo. ^23^ The stiffness of the ECM is another factor that influences cellular metabolism, migration, and arrangement in a tissue. ^24^ In our study, before culturing the PLSM cells in the chip we optimised the hydrogel concentration to ensure the even distribution of cells inside the hydrogel and appropriate cell-cell interaction inside the 3D environment.

We used three different concentrations of Rat tail Collagen 1 to culture PLSM cells (1 mg/mL, 1.5 mg/mL, and 2 mg/mL) to confirm the most desired hydrogel concentration (Fig. S1). Live/dead staining using calcein-AM (Fig. 1a) and PI (Fig. 1b) staining showed that 2mg/mL of collagen 1 supports even distribution of the cells (Fig. 1d) and it was chosen for all the experiments inside the chip. This will ensure the close interaction between cells in the 3D environment.

**Fig 1.**
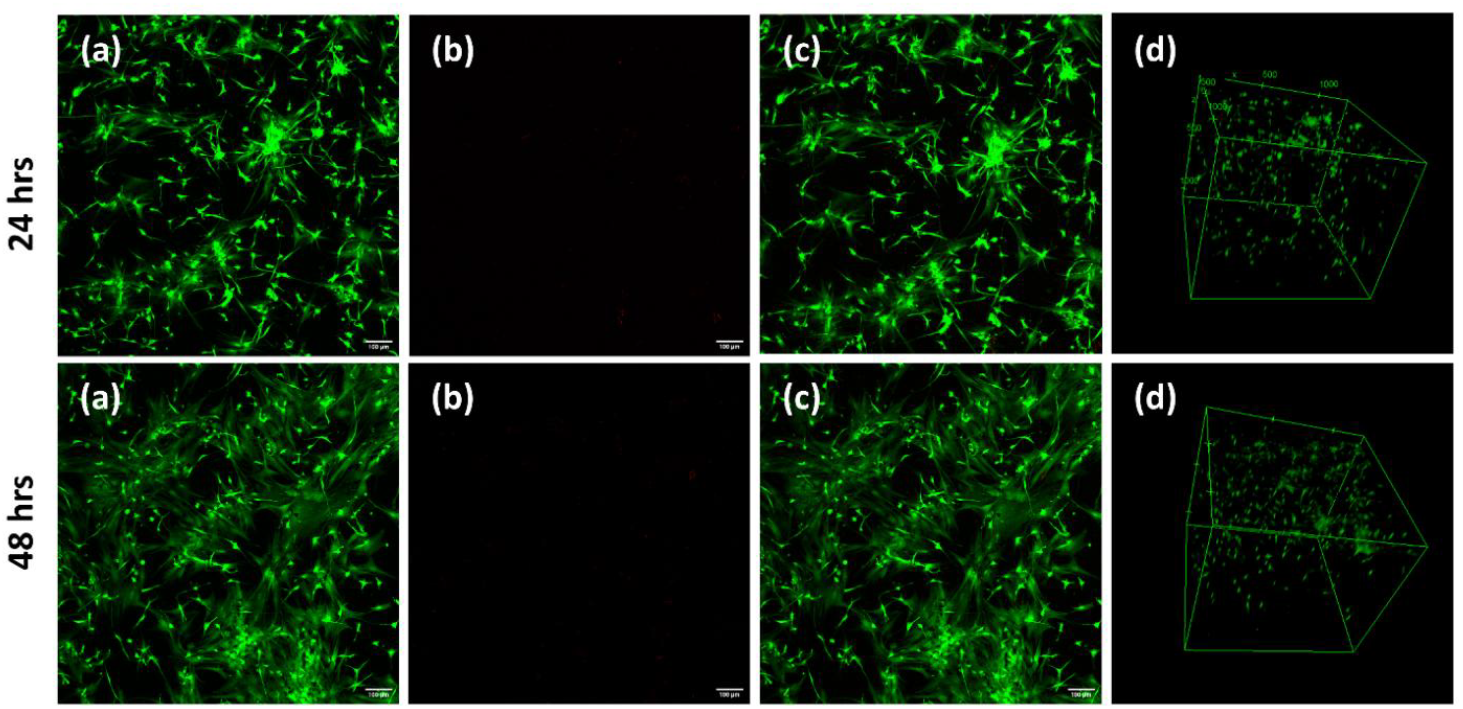
Gel optimization (collagen-2mg/mL) by live dead assay using Calcein AM and PI staining in confocal laser scanning microscope after 24 hrs and 48hrs; (a) Calcein AM, (b)Propidium iodide, (c) combined image, (d) 3D view (magnification 10x, error bar 100µm

### Design, fabrication of the chip

ASM microfluidic chip consists of three channels separated by two arrays of circular pillars. This provides a central channel for holding the hydrogel and two side channels on either side for media perfusion, each having separate inlets and outlets (Fig 2a). The chip is fabricated in a single layer of PDMS bonded onto a coverslip using plasma treatment (Fig 2c), this bonding to the coverslip allows for higher magnification imaging under the confocal microscope. Microfluidic devices with ‘parallel channel design’ features micropillars of different sizes and shapes ^25–29^ for gel-holding compartments. In the present study, the pillars are circular which is easier to fabricate while considering micro milling for device mould preparation (Fig 2b). The circular pillars can effectively hold the hydrogel when loaded manually using a pipette. When the gel is loaded via the inlet of the central channel, it will hold in place without leakage unless the pressure inside the gel chamber is lower than the pressure between the pillars. This was achieved by pipetting the gel slowly and steadily at an approximate flow rate of 10 µl per 20 seconds (Fig 2d). Once the hydrogel is set the side channels can be used to supply nutrient media and other chemicals to the hydrogel.

**Fig 2.**
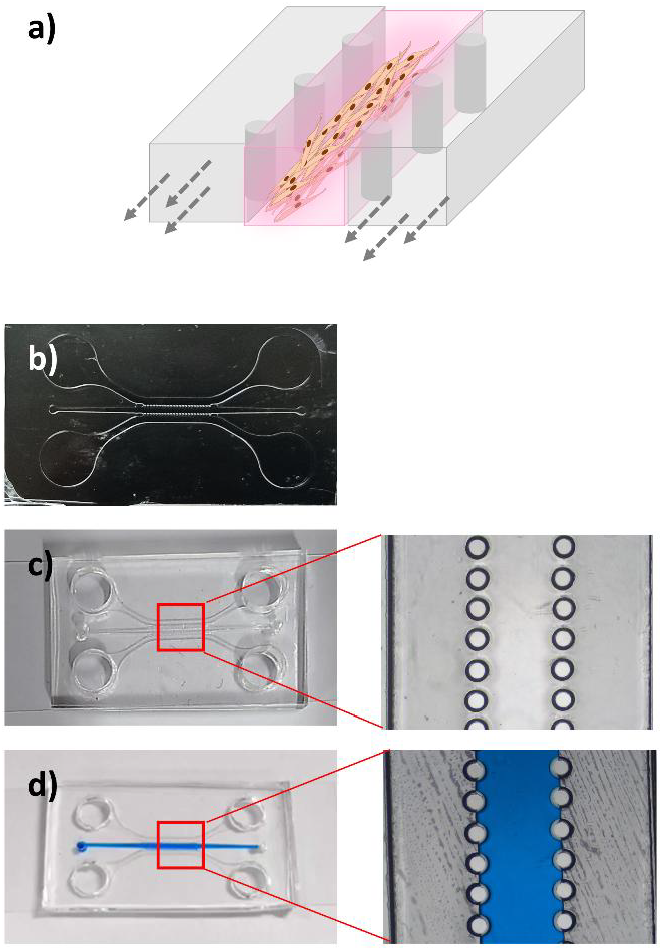
(a) Schematics of the chip (b)PMMA mould fabricated using CNC milling (c) PDMS chips fabricated using replica moulding (d) Chip characterisation using coloured media.

ASM cells are embedded beyond alveolar epithelial and fibroblast cells in human lungs, therefore the signalling molecules like inflammatory cytokines and other growth factors reach ASM cells via diffusion. The architecture of the chip allows manipulation of the hydrogel without disturbing the embedded cells by perfusing necessary compounds via the side channel. During a lung injury, epithelial cells are directly exposed to the xenobiotic compounds however the local and systemic biological response is mediated by all surrounding cells including lung fibroblast, lung smooth muscle cells and immune cells. By the release of proinflammatory cytokines at the site of exposure, the signalling molecules recruit immune cells and signal the fibroblasts and smooth muscle cells to undergo necessary changes like repair, fibrosis and remodelling. This flow of signals is mediated by the diffusion of specific molecules through the ECM at the site. Repeated injury and repair in the alveolar tissue leads to permanent remodelling of the airway. There is now considerable evidence that suggests ASM has a crucial role in this pathophysiology of respiratory diseases like asthma, by controlling the ECM production, cell proliferation, migration and changing the contractile function. ^30^ Hence modelling an ASM in its 3D microenvironment is crucial to understanding the mechanisms underlying the remodelling of the airway in different pathological conditions. Our chip provides this feature with ease, the PLSM cells are embedded in a 3D hydrogel which can be stimulated with any signalling molecules.

### Numerical simulation

The design of the microfluidic chip consists of a central gel channel, flanked by two side channels, with an array of micropillars that serve to demarcate the central channel from the adjacent side channels. In the mesh convergence study, we observed as the number of meshing elements decreased, the relative errors in collagen solution filling increased. Consequently, the simulation study was done using fine mesh (13862 triangular elements) with an average element quality of 0.8126 (Fig S2). The successful filling of the gel within the central channel hinges on meticulously balancing capillary forces against surface tension. The hydrophobic surfaces allowed the gel or fluid to be filled without leakage rather than hydrophilic surfaces ^31^. Another study by Lee et al. presented a computational fluid dynamic simulation for the sequential gel filling process within the channels ^32^. A study by Ghobadi et al. identified that micropillar gap spacing up to 0.1mm is effective in containing gels within a 0.9 mm wide channel. Consistent with previous studies, this study successfully simulated the gel-filling process (Fig 3). The experimental validation using blue-coloured water and collagen for the gel filling is shown in Figure 2d.

**Fig 3.**
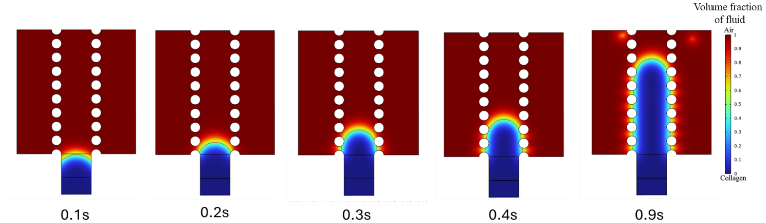
Simulation of hydrogel filling into the microfluidic device at different time intervals; Movement of collagen without leakage along the central gel channel of the design is simulated; The dynamic viscosity of the 2mg/mL collagen solution is 6mPa.sec.

### ASM on chip characterisation

In 2D culture flasks PLSM cells grow as individual cells attached to the flak bottom. Inside a 3D matrix, it could show the contractility as the hydrogel is soft and shrinkable. We loaded the hydrogel embedded with a high density of PLSM cells inside the gel-holding compartment of the chip and allowed for gelation. The chip was supplemented with nutrient media and was cultured for 5 days initially. Brightfield images of day 0 to 5 shows remarkable cellular orientation of PLSM cells inside the chip. After 24 hours inside the chip, cells are observed to pull the hydrogel and start to form a single fibre-like structure. By day 5, individual cells become undistinguishable and form a thick band of fibre in the middle of the channel (Fig 4). This self-orientation of the PLSM cells is understood to be cell-dependent reorientation, as the alveolar epithelial cell line (A549 cells) seeded inside the chip failed to reorganise into any kind of structures observed in PLSM cells (Fig S3). This might be because of the contractile property of the PLSM cells. This finding indicates that cells are forming fibre-like structures because of their contractile property, not because of the design of the chip.

**Fig 4.**
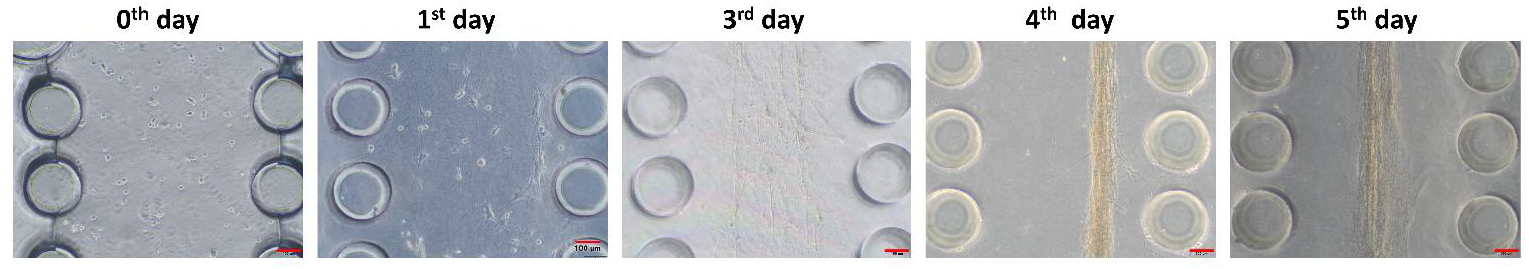
Cell culture optimisation (collagen-2mg/ml), 3D cultures of primary lung smooth muscle cells in microfluidic devices, images were captured using Olympus Trinocular microscope, 10x objective, scale bar=100µm

To characterise the PLSM cells and the changes they acquired inside the chip, we performed Live/dead assay by calcein-AM and PI staining and studied the arrangement of the cytoskeleton by Rhodamine-phalloidin staining and the cellular orientation by DAPI staining.

From the Live/dead assay, we observed that cells are viable even after 7 days of culture inside the chip with daily media supplementation (Fig 5a). This sparks a great step towards considering microfluidic chips for long-term toxicity studies that are necessary for regulatory approvals. The Rhodamine-phalloidin staining of F-actin shows the longitudinal arrangement of the cytoskeleton in the muscle fibre formed (Fig 5b). In elongated cells like PLSM, nuclear orientation shows the orientation of the cell. From DAPI-stained images, nuclear orientation was calculated using the image analysis technique. From the images taken on days 2 and 7, data shows that an increasing number of cells orient themselves towards a single direction as the days progress. This indicates that the cells are dividing and aligning in the same direction as the fibre alignment (Fig 5c).

**Fig 5.**
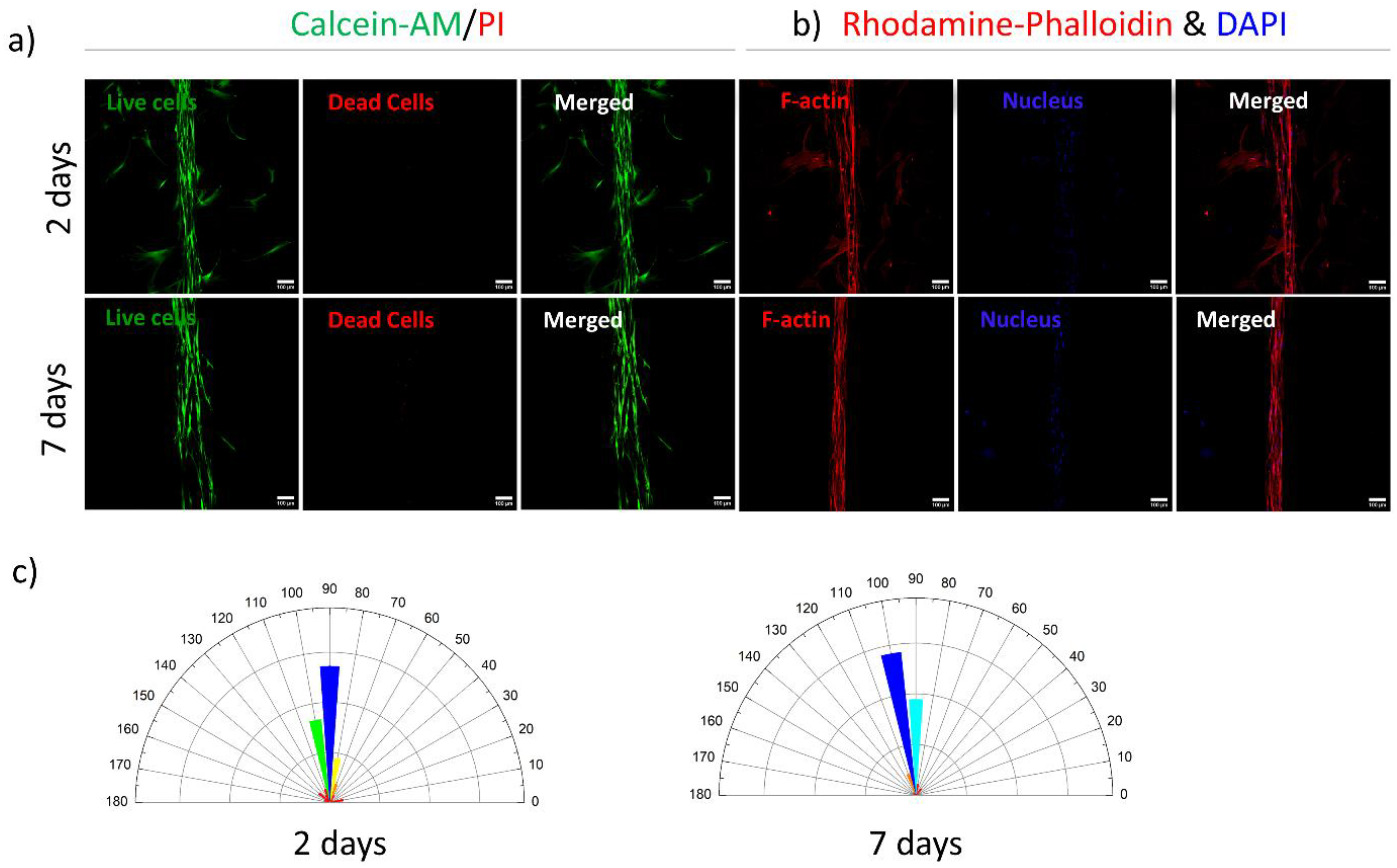
(a) Live/Dead staining using Calcein AM/PI double staining after 2 days and 7 days, under confocal laser scanning microscope; max imum intensity projection of stacked image magnification 10x, scale bar-100µm (b), Cytoskeleton imaging using Rhodamine phalloidin and counterstain using DAPI after 2 days and 7days, under confocal laser scanning microscope; maximum intensity projection of stacked image magnification 10x, scale bar-100µm (c) rose diagram representing nuclear orientation.

The fabricated chip allows the indirect exposure of molecules to the cells via diffusion. Diffusion of 70kDa FITC-Dextran was visualised in the chip under the confocal microscope to confirm the movement of molecules from the side channels to the central channel. The FITC-Dextran molecules get diffused into the central channels in under 30 minutes (Fig S4).

### Effect of TGF-β on ASM remodelling

ASM cells have phenotypic plasticity, meaning they can change their behaviour and function to a degree. Two major phenotypes of ASM cells are the contractile and synthetic or proliferative phenotype. They are speculated to coexist in a balance inside our body. Imbalance in either population leads to pathological conditions like asthma and COPD. ^33^ Several intrinsic and extrinsic factors affect the phenotypic change in AMS cells. These reversible switches are called ASM remodelling. ECM proteins (Collagen, fibronectin, laminin etc.), Growth factors (Platelet Derived Growth factor, TGF-β etc.), and inflammatory cytokines (IL-4, IL-13 and TNFα etc.) are some of the intrinsic factors that can initiate a phenotypic change in ASM cells. ^34^ This release of signalling molecules can be due to extrinsic factors like inhalation of toxicants, smoke and other infections. To understand the effects of a signalling molecule on ASM we exposed TGF-β via the side channels. Upon exposure for 7 days, the effects were studied using measuring the gel compaction, cell number and expression of contractile protein calponin 1 and structural protein F-actin. We observed a significant difference between the percentage reduction in hydrogel width from day 0 to day 7, with more reduction in the TGF-β-group than the TGF-β+ group (Fig 6a-b). we also observed an increase in the number of cells in the TGF-β+ group after 7 days compared to the TGF-β-group (Fig 6c). This change is expected since pro-inflammatory mediators like TGF-β are known to increase ECM secretion, cell size and number. ^11,35^ TGF-β is a growth factor that is produced as a pro-inflammatory mediator by fibroblasts under different pathological conditions. The release of TGF-β and other pro-inflammatory cytokines induce excessive ECM secretion like collagen and fibronectin. In ASM cells, TGF-β acts as an autocrine and paracrine signalling molecule that initiates differentiation and proliferation and increases the ASM cell size. ^11,30,36^We observed no significant difference between the expression of F-actin (Fig 7a-b) and Calponin-1 (Fig 7c-d in the two groups except in the case of F-actin at day 2, F-actin in the TGF-β-group was significantly higher than the TGF-β+ group. Expression of both F-actin and Calponin 1 protein is reported to be high in ASM cells in contractile phenotype. ^37,38^ There are other factors that contribute to the phenotypic change in ASM cells that might have influenced the results, like serum deprivation, the presence of collagen in the hydrogel etc.

**Fig 6.**
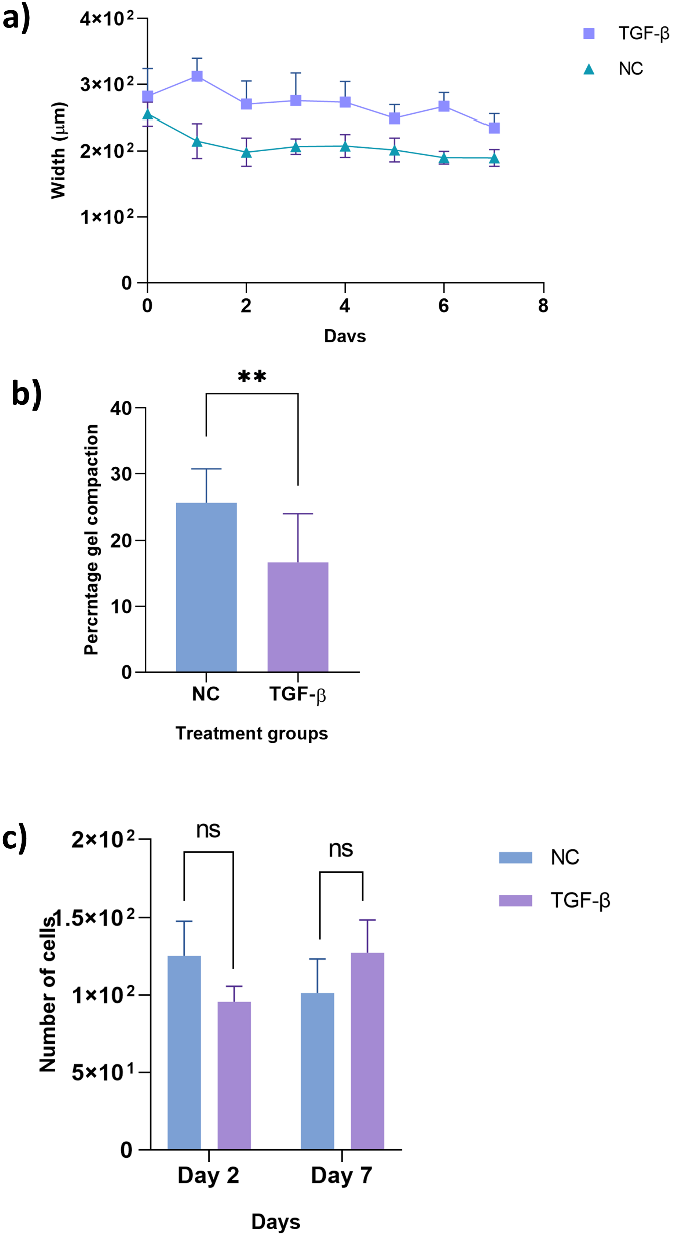
(a) Changes in the width of the gel channel due to the TGF-β treatment (b) Gel compaction after 7 days in ASM-on-a-chip exposed to TGF-β, (c) Changes in the cell number.

**Fig 7.**
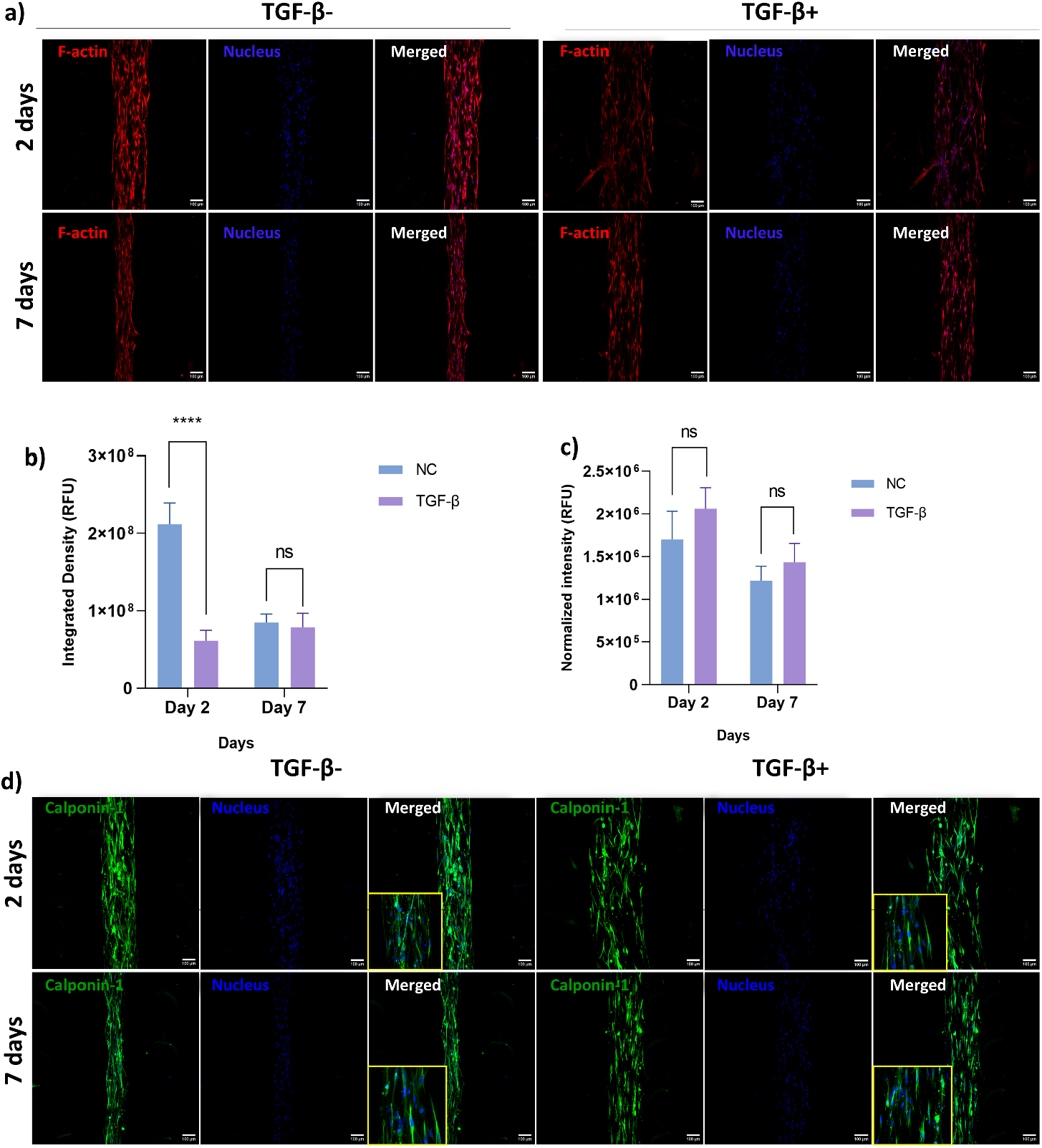
(a) Rhodamine phalloidin staining of F-actin after the exposure to TGF-β (b Quantification of F-actin intensity from cytoskeleton imaging after 2 days and 7day due to the treatment of TGF β (c) Changes in the normalised fluorescent intensity of calponin-1 due to the exposure of TGF β (d) Immunofluorescence staining of Calponin-1 after the exposure to TGF β. Images taken under confocal laser scanning microscope; maximum intensity projection of stacked image magnification 10x, scale bar-100µm, magnified 20x images are added as inserts.

## Conclusion

The lung is an injury-prone organ, and lung cells are exposed to numerous particles in the atmosphere abundantly. The consequences of such alveolar insults lead to acute and chronic inflammation and infections. This could potentially lead to respiratory disorders like asthma and COPD. Studies have found that the manifestation of these respiratory disease symptoms is due to a combined effect of epithelial cells, fibroblast cells and ASM cells. The insult-derived responses of epithelial cells and fibroblasts release numerous signalling molecules that create local inflammation and remodelling. The role of ASM cells in manifesting symptoms is reported and as ASM remodelling is a reversible process, therapeutics that enable re-remodelling of ASM would be worth researching for. While literature reports conclusive effects of TGF-β like inflammatory molecules on ASM remodelling, the lack of robust and versatile 3D models of ASM has been a major gap in this field. Cofounding factors that influence the ASM remodelling in vitro models like serum conditions and the presence of ECM proteins in the hydrogel might create disparity in the obtained result, however, the 3D model of ASM for studying remodelling is well established by our ASM-on-chip device. In the present study we have established that ASM-on-chip can be used for long-term cultures and to model ASM functions in normal and pathological conditions. The microfluidic design provides easy access to ASM cells that are embedded in the hydrogel via diffusion, this recapitulation of in vivo conditions further enhances the potential applications and the relevance of the chip. The circular micropillars are shown to effectively hold the gel in place as the other shaped micropillars, which highlights the simplicity and ease of fabrication of ASM-on-chip compared to other micropillar structures. The chip also provides a long-term window of observation on how smooth muscle cells self-orient themselves inside the matrix in real-time, this is achieved for the first time in a microfluidic device. The role of ECM on tissue remodelling is another important aspect that can be extensively investigated using the ASM-on-chip. This study was carrier out with collagen 1 hydrogel, however by altering the composition and viscoelasticity of ECMs studies can be taken further on the front of effects of ECM on tissue remodelling. The growth of reoriented ASM cells parallel to the channels enables an extensive cell-cell and cell-ECM communication interfaces, and with the possibility two interfaces on both sides this virtue is enhanced and boost the potential applications of the chip. ASM-on chip can be used as a new alternative platform that covers different aspects about common respiratory diseases like asthma and COPD, and lung injuries caused due to inhalation of hazardous particles (air pollutants, vapes, particulate matters etc.) as mentioned above. Furthermore, the possibility of adding more cells like epithelial cells and fibroblast cells in side channels to see how cellular cross-talk determine disease progression or revival, and the possibility of introducing mechanics via continuous perfusion through side channels projects ASM-on-chip as a potent replacement model for airway diseases.

## Author contributions

(AS) Conceptualization, Data curation, Investigation, Methodology, Visualization, Writing – original draft; (JX) Conceptualization, Data curation, Investigation, Methodology, Visualization, Writing – original draft, review& editing; (HPK) Investigation, Methodology, Writing – original draft; (AK)-Investigation, Software, Visualization; (AP) Investigation, Methodology, Visualization; (MKB) Investigation, Methodology, Visualization;(RS) Investigation, Methodology, Visualization; (RNS) Funding acquisition, Resources, Supervision; (JBS) Conceptualization, Data curation, Investigation, Writing – review& editing, Funding acquisition, Resources, Supervision.

## Data availability

The data supporting this article have been included as part of the ESI.

## Acknowledgements

AS, AK and JX acknowledge the support from CSIR, Govt. of India for the SRF Fellowships. RS JX, JBS acknowledges funding from the UK Foreign, Commonwealth & Development Office (FCDO) for the CSC split-site fellowship (INCN 2021-143). RS thanks UGC, Govt. of. India for the SRF fellowship. J.B.d.l.S acknowledges support from BBSRC (BB/V019791/1) and the Integrated Biological Imaging Network (IBIN), a Technology Touching Life MRC Network (MR/W024985/1). Also funding from MRC (MR/X013855/1), and the Wellcome Trust (301619/Z/23/Z).

