## Supplementary figures and images for "Airway smooth muscle--on-a-chip: a microfluidic approach to study alveolar smooth muscle remodelling"

### Fig S1

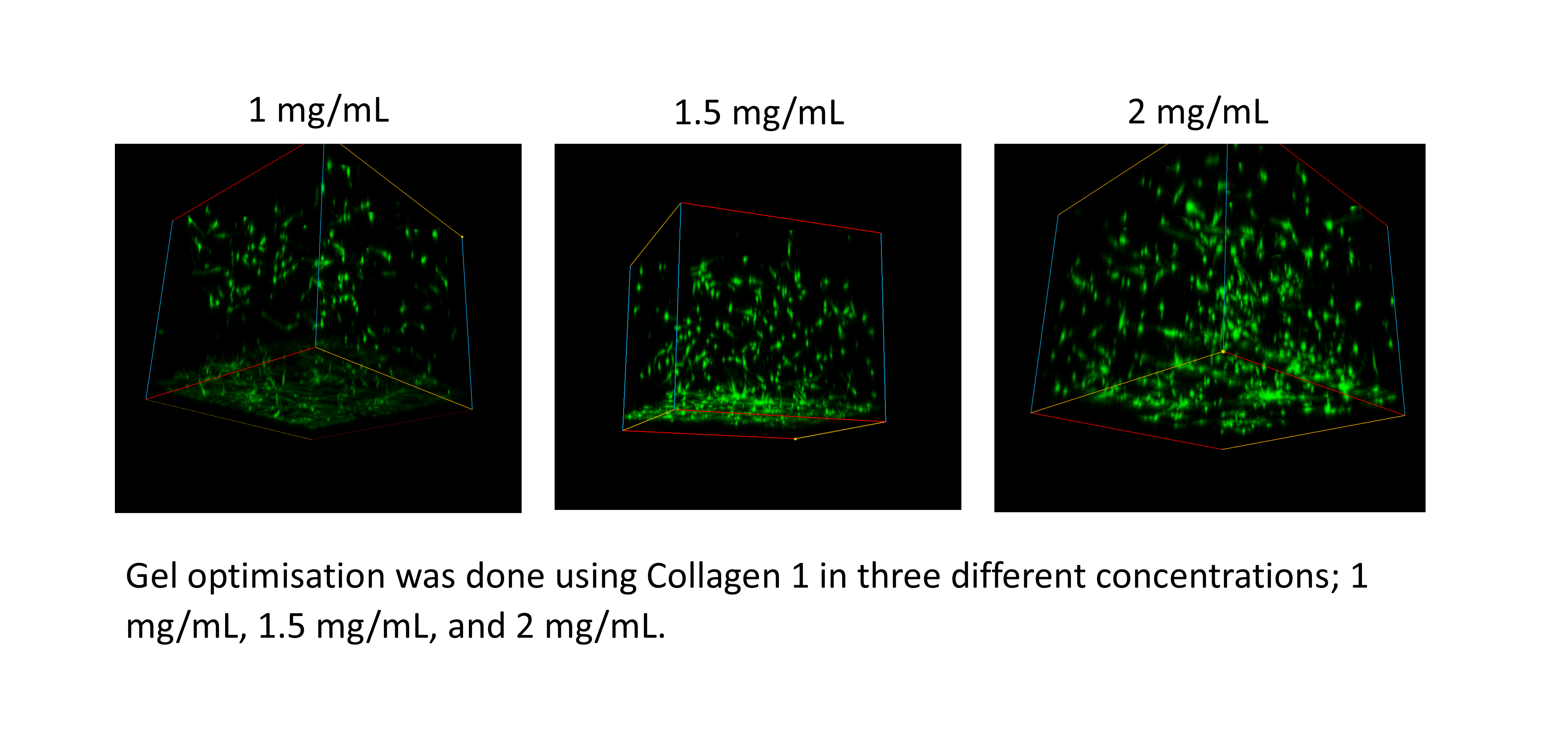

### Fig S2

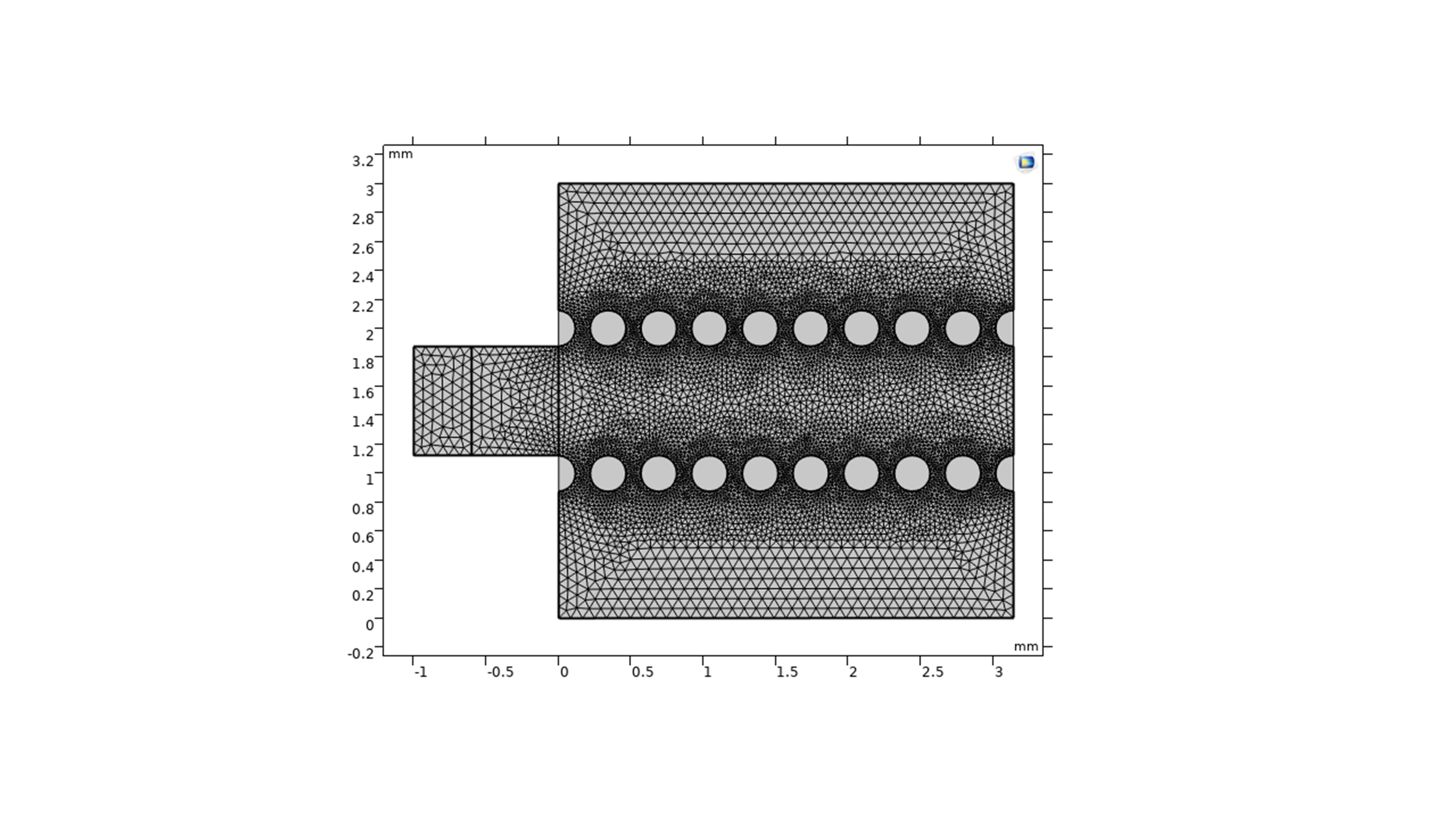

### Fig S4

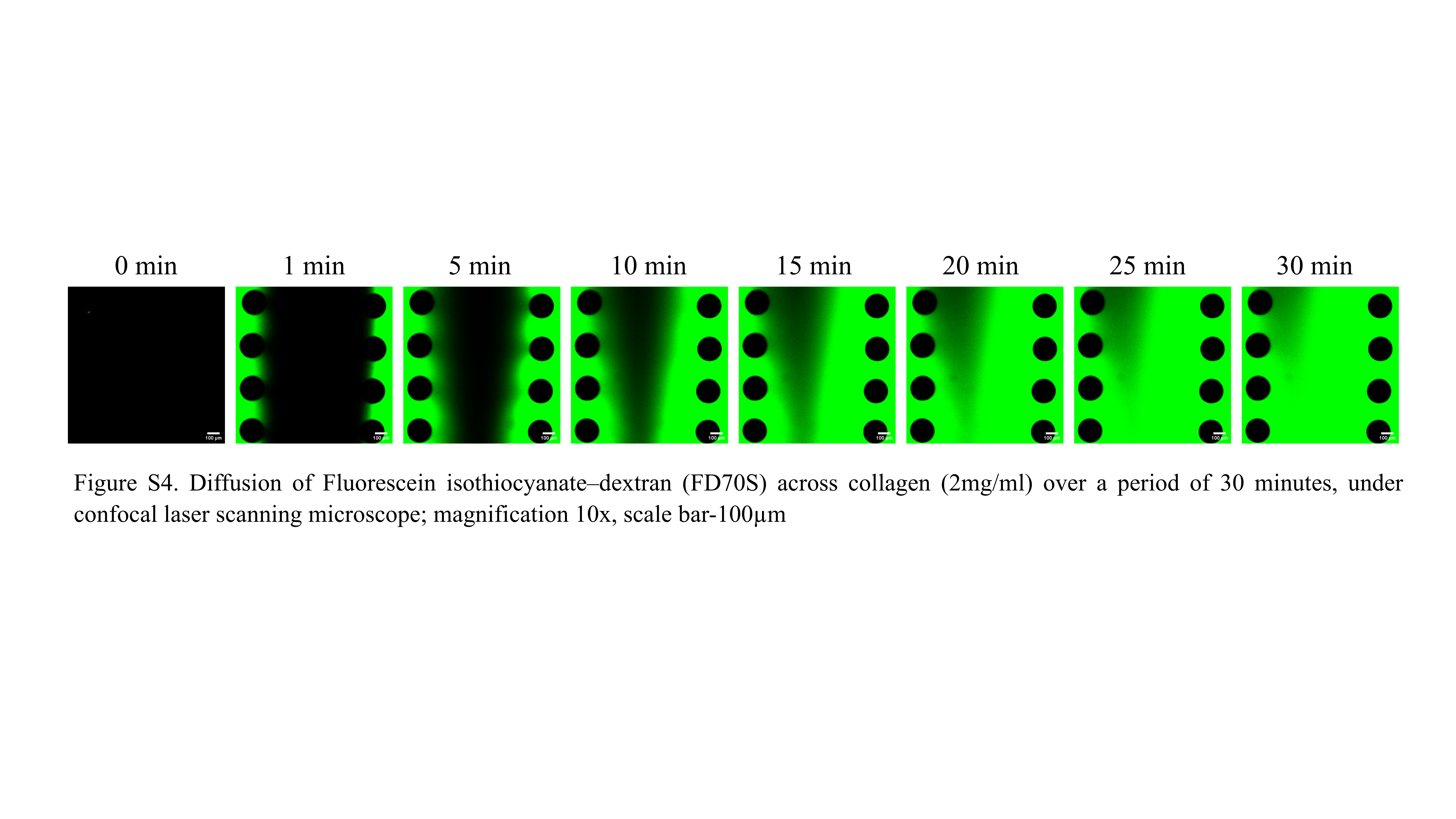

### Fig S43

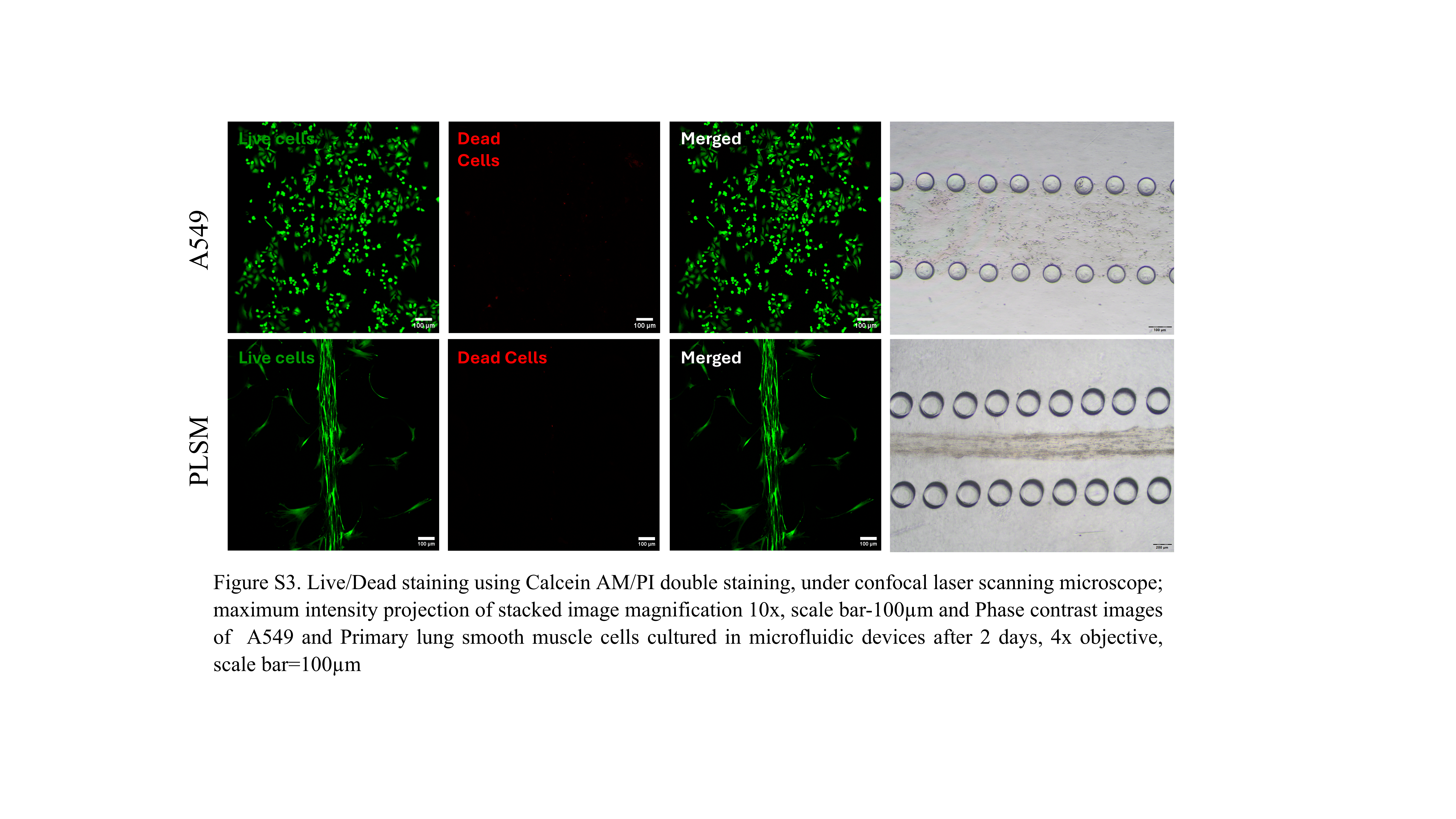
